# Prototyping a valinomycin biosynthesis pathway within a cell-free transcription-translation (TX-TL) system

**DOI:** 10.1101/091520

**Authors:** Tiffany Zhou

## Abstract

Many natural metabolites have antibacterial, antiviral, or anticancer effects and can be developed into new drugs. However, working with the microorganisms that produce these products can be challenging since they are not as well characterized as a model organism like *Escherichia coli*. In this paper, we investigate the potential for a cell-free transcription-translation (TX-TL) system to provide a rapid prototyping platform for characterizing new genetic pathways. We use the valinomycin biosynthesis pathway as a test case, and we show successful heterologous expression of the heterodimeric valinomycin synthetase (VlmSyn, Vlm1: 374 kDa and Vlm2: 284 kDa) from *Streptomyces tsusimaensis* within the TX-TL system. Using LC-MS analysis, we find that valinomycin is produced at low but detectable levels, even when only one out of the three basic precursors is fed into the system. Our work represents another step towards applying cell-free biosynthesis to the discovery and characterization of new natural products.

## 2 Introduction

Natural products have been a key source for new drugs, including antibacterial, antiviral, and anticancer compounds, for over 30 years [1]. Recent developments in large-scale genetics and metabolomics have revealed that the chemical space of secondary metabolites still remains largely unexplored [2]. However, the characterization of new biosynthetic pathways and the small molecules they produce has been difficult, partly due to the challenges of working with the natural product producing organisms in the laboratory. Synthetic biology can solve this problem through robust heterologous expression of the biosynthetic pathways within more well-understood host organisms, such as *Escherichia coli* or *Saccharomyces cerevisiae*.

Cell-free protein expression systems are quickly emerging as a useful tool for synthetic biology research. They take advantage of native transcription-translation machinery and allow for the simultaneous *in vitro* expression of multiple genes in the form of either plasmid or linear DNA [3,4]. Cell-free systems offer many benefits over cell-based expression, including the elimination of the need for an appropriate host strain, the ability to easily control specific reaction conditions, the absence of cell processes that could extraneously influence the behavior of the targeted gene network, and a greater tolerance for proteins and metabolites toxic to living cells [3]. Reactions in cell-free systems can be performed in a day, as opposed to one week with *in vivo* methods, allowing for more rapid design-build-test cycles [5]. Researchers have recently shown cell-free systems to have potential in the areas of rapid pathway prototyping [3–5], genetic network design [3,6], and high-yield protein production [7].

To investigate the capability of cell-free systems for the characterization of heterologous natural biosynthetic pathways, we decided to test the biosynthesis pathway for valinomycin in an *E. coli*-based cell-free transcription-translation (TX-TL) system developed by Noireaux [8]. Valinomycin is a dodecadepsipeptide produced by various *Streptomyces* strains that exhibits a range of antibiotic, antifungal, and antiparasitic activities [9]. A gene cluster capable of synthesizing valinomycin has been identified, and it is proposed that its biosynthesis involves a nonribosomal peptide synthetase (NRPS) called tetramodular valinomycin synthetase (VlmSyn) [10]. NRPSs are challenging to express in heterologous platforms because of their large sizes (typically ranging from 100 to *>*300 kDa) and complex structures, but not impossible since two NRPS proteins GrsA and GrsB were previously reported to show high expression and functionality within an *E. coli*-based cell-free system [11]. In addition, researchers were successful in heterologously expressing VlmSyn from *Streptomyces tsusimaensis* in a soluble and active form in *E. coli* [9]. It was found that the *in vivo* production of valinomycin in *E. coli* could occur without precursor feeding or optimization of cultivation conditions.

VlmSyn is encoded by two distinct genes *vlm1* (10,287 bp) and *vlm2* (7,968 bp) [9], and they form a protein “assembly line” of various functional domains organized into four modules (Figure 1). The process requires three basic precursors: *α*-ketoisovalerate (*α*-Kiv), pyruvate (Pyr), and L-Valine (L-Val). *α*-Kiv is reduced to D-*α*-hydroxyisovaleric acid (D-Hiv) via the ketoreductase (KR) domain in Module 1. The epimerase (E) domain in Module 2 transfers L-Val to D-Valine (D-Val). The KR domain in Module 3 reduces pyruvate to L-Lactate (L-Lac). These three products (D-Hiv, L-Lac, and D-Val), along with L-Val, are linked to form a tetradepsipeptide basic unit. A terminal thioesterase (TE) domain oligomerizes and cyclizes the repeating sequence of this basic unit to form the final valinomycin product [10].

In this report, we demonstrate successful expression of VlmSyn in the TX-TL cell-free system and show that VlmSyn produces detectable amounts of valinomycin in TX-TL, even when only one out of the three biosynthetic precursors is fed into the system.

**Figure 1:**
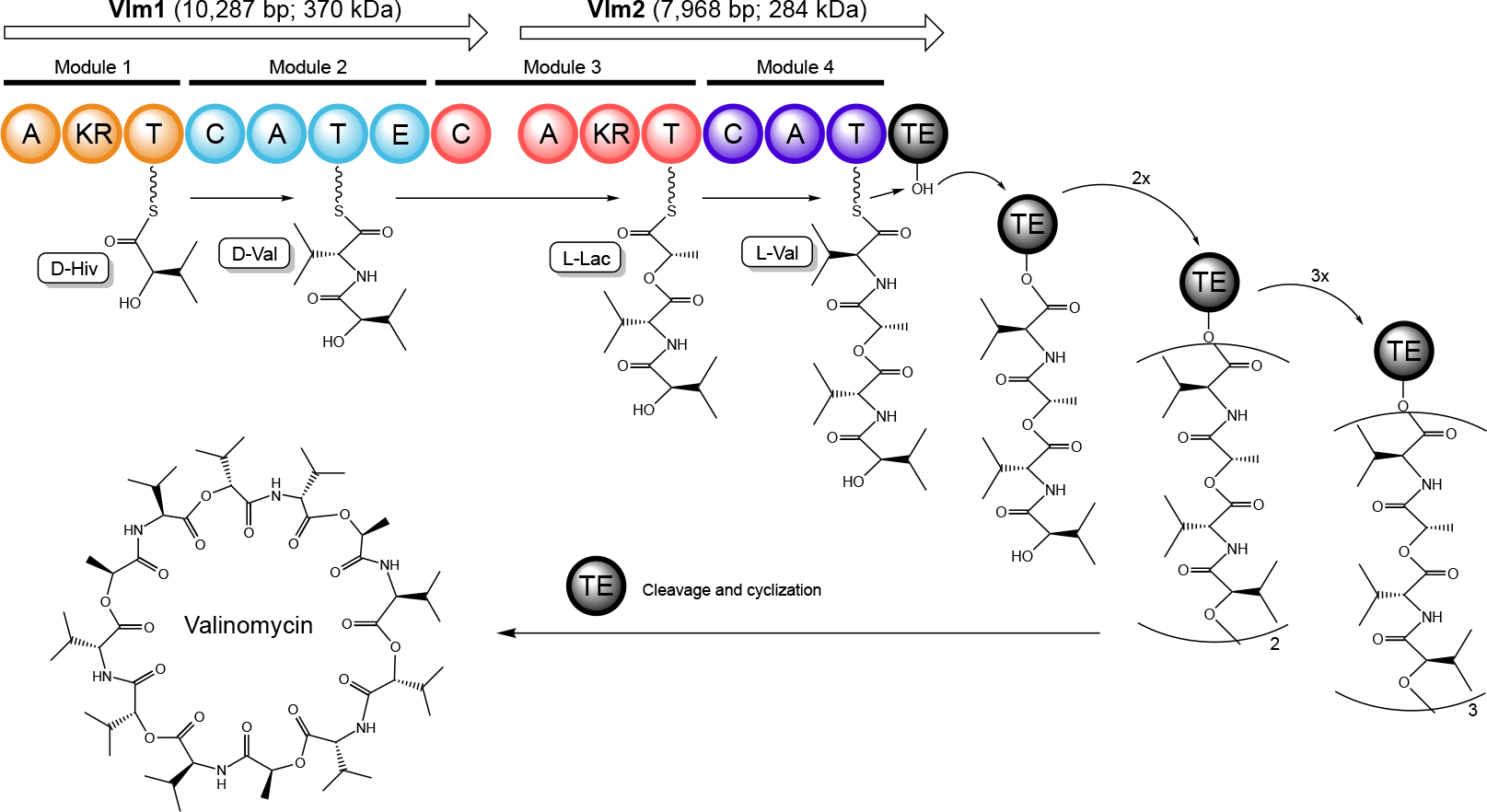
Proposed model for the biosynthesis of valinomycin. The valinomycin synthetase is divided into four modules, consisting of domains with adenylation (A), ketoreductase (KR), thiolation (T), condensation (C), epimerase (E), and thioesterase (TE) functions. Each module incorporates D-Hiv, D-Val, L-Lac, or L-Val to form a tetradepsipeptide basic unit. After three units are formed, they undergo cleavage and cyclization to form the final valinomycin product. (Figure inspired by models in [9], [10].)

## 3 Results/Discussion

### 3.1 Obtaining *vlm1* and *vlm2* genes

We obtained three plasmids that Jaitzig, et al. [9] constructed during their experiments to reconstitute valinomycin biosynthesis in *E. coli* (Table S1, Supplementary Material). The first two plasmids, pVlm1 and pVlm2, contain the *vlm1* gene and the *vlm2* gene, respectively, controlled by a *plac* promoter. As a side experiment, we tried to extract *vlm1* and *vlm2* from these plasmids using a PCR protocol modified to account for large amplicon lengths. The modifications included increasing the template concentration, lengthening the initial denaturation time, increasing the number of cycles, and lengthening the extension time and extension temperature. In addition, we tested a gradient of annealing temperatures between 68°C and 75°C. Gel electrophoresis showed strong bands at the correct lengths for *vlm1* and *vlm2* (Figure 2). However, we also saw multiple nonspecific bands, especially at lower annealing temperatures, which could have resulted from non-target primer binding or primer-dimer formation. Further tests to determine the optimal PCR conditions are needed in order to obtain *vlm1* and *vlm2* in linear form.

The third plasmid, pET15b-sfp, contains the *sfp* gene controlled by a T7 promoter. *sfp* encodes a phosphopantetheinyl transferase (PPTase) enzyme Sfp from *Bacillus subtilis* that has been shown to function in numerous heterologous organisms [9,12]. PPTases are needed to transfer a phosphopantetheine group during a post-translational modification step for NRPSs like VlmSyn. Even though *E. coli* contains native PPTases, it was shown that Sfp is more effective at accepting Vlm1 and Vlm2 as substrates [9].

**Figure 2:**
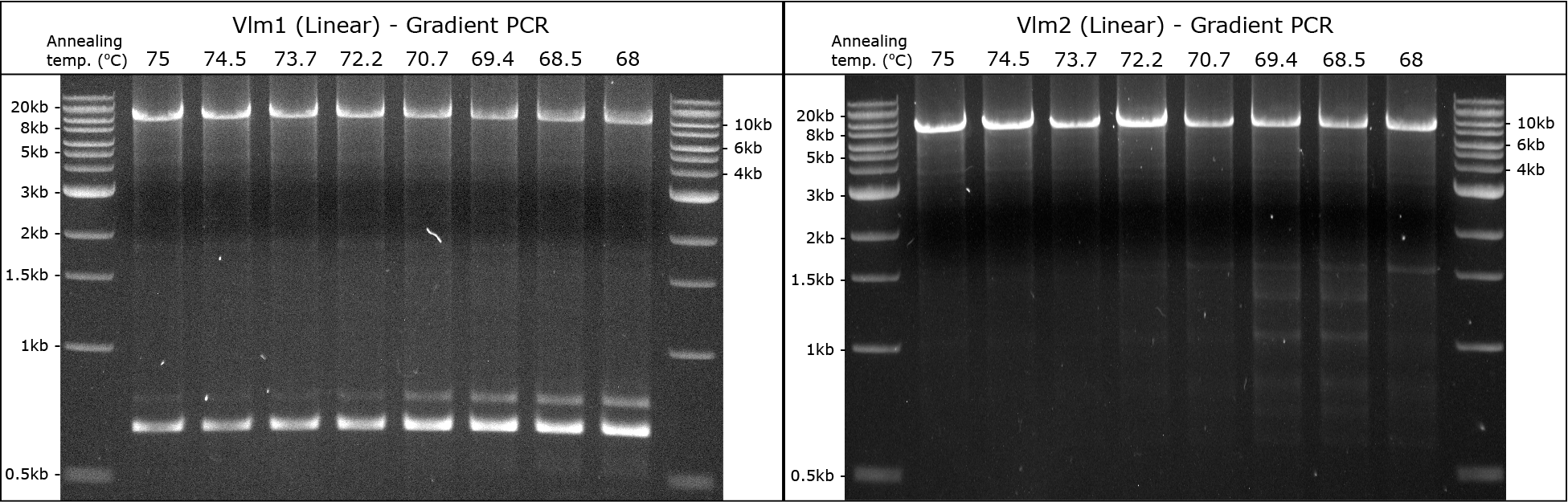
Gel electrophoresis analysis for PCR of *vlm1* (left) and *vlm2* (right). A gradient of annealing temperatures between 68°C and 75°C was tested. Expected sizes for *vlm1* and *vlm2* linear products are 10,287 bp and 7,968 bp, respectively.

### 3.2 Creating TX-TL extract from *E. coli* BJJ01

In order for valinomycin biosynthesis in TX-TL to work, we need to express the PPTase Sfp in the system. One way to do this is to add the *sfp* plasmid to every TX-TL reaction, meaning that each reaction would have three different plasmids. However, resource limitations in TX-TL could prevent simultaneous expression of all three plasmids [13], especially since the *vlm1* and *vlm2* genes are so large. To avoid this problem, we decided to create TX-TL extract directly from *E. coli* strain BJJ01, which Jaitzig, et al. created by integrating the gene *sfp* under the control of a constitutive promoter into the genome of *E. coli* BL21 Gold.

Since this strain constitutively produces Sfp, the enzymes will already be present in its TX-TL extract. We created the extract following the methods described by Sun, et al. [14], with the exception that we used homogenization instead of bead-beating during the cell lysis step. The buffer calibration results (Figure 3) show that this new extract (which we named “E34”) requires a buffer containing 4mM of Mg and 280mM of K, and that its GFP expression can reach over 5000 nM.

**Figure 3:**
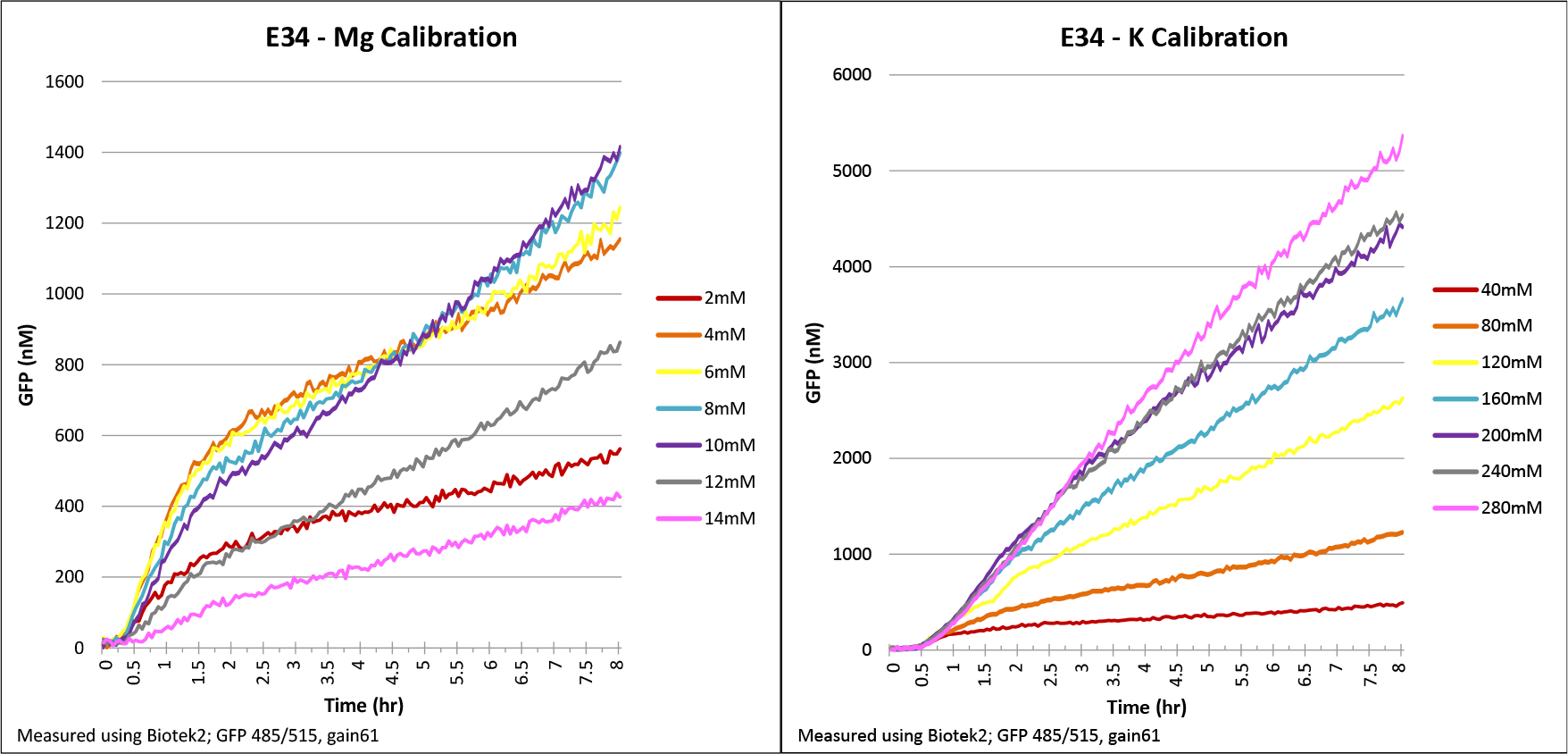
Buffer calibration results for E34 extract, which was made from *E. coli* strain BJJ01. Concentrations for the final buffer are 4mM of Mg and 280mM of K.

### 3.3 Expressing Vlm1 and Vlm2 in TX-TL

To test Vlm1 and Vlm2 expression in our new E34 extract, we first added pVlm1 and pVlm2 with concentrations varying between 0.001nM and 10nM into separate TX-TL reactions. We used 20nM of IPTG to induce expression and ran the reactions at 25°C for 12 hours. We decided to use a temperature lower than the usual 29°C because it slows down translation, which avoids potential protein folding issues for the large enzymes. Using SDS-PAGE to assess protein expression, we found that a large protein above 200kDa is expressed when plasmid concentrations are 1nM or higher (Figure 4). These are very likely to be our target VlmSyn proteins, since no proteins of that size are found in the TX-TL extract itself. A Western blot can be performed in order to further confirm Vlm1 and Vlm2 expression.

Next, we held the plasmid concentrations constant at 1nM and varied the IPTG concentration and the reaction temperature to find the optimal reaction conditions. Once again, Vlm1 and Vlm2 expression were tested separately, and the reactions were run for 12 hours. The SDS-PAGE results (Figure 5) show that varying the reaction temperature affects expression more than varying the IPTG concentration. In fact, as the temperature increases, multiple bands above 200kDa begin to appear. We believe that since translation speeds up at higher temperatures, errors could occur during the production of the large proteins, leading to truncated or incomplete sequences that show up in SDS-PAGE. It may be useful to move the His-tag to the C-terminal and run a Western blot to confirm whether complete VlmSyn proteins are being produced.

We also see large-size bands in our negative controls (0nM IPTG), which was not expected. However, these controls do contain 1nM of the Vlm1 or Vlm2 plasmid, so the presence of the large bands even in the absence of IPTG suggests that the *plac* promoters could be leaky.

**Figure 4:**
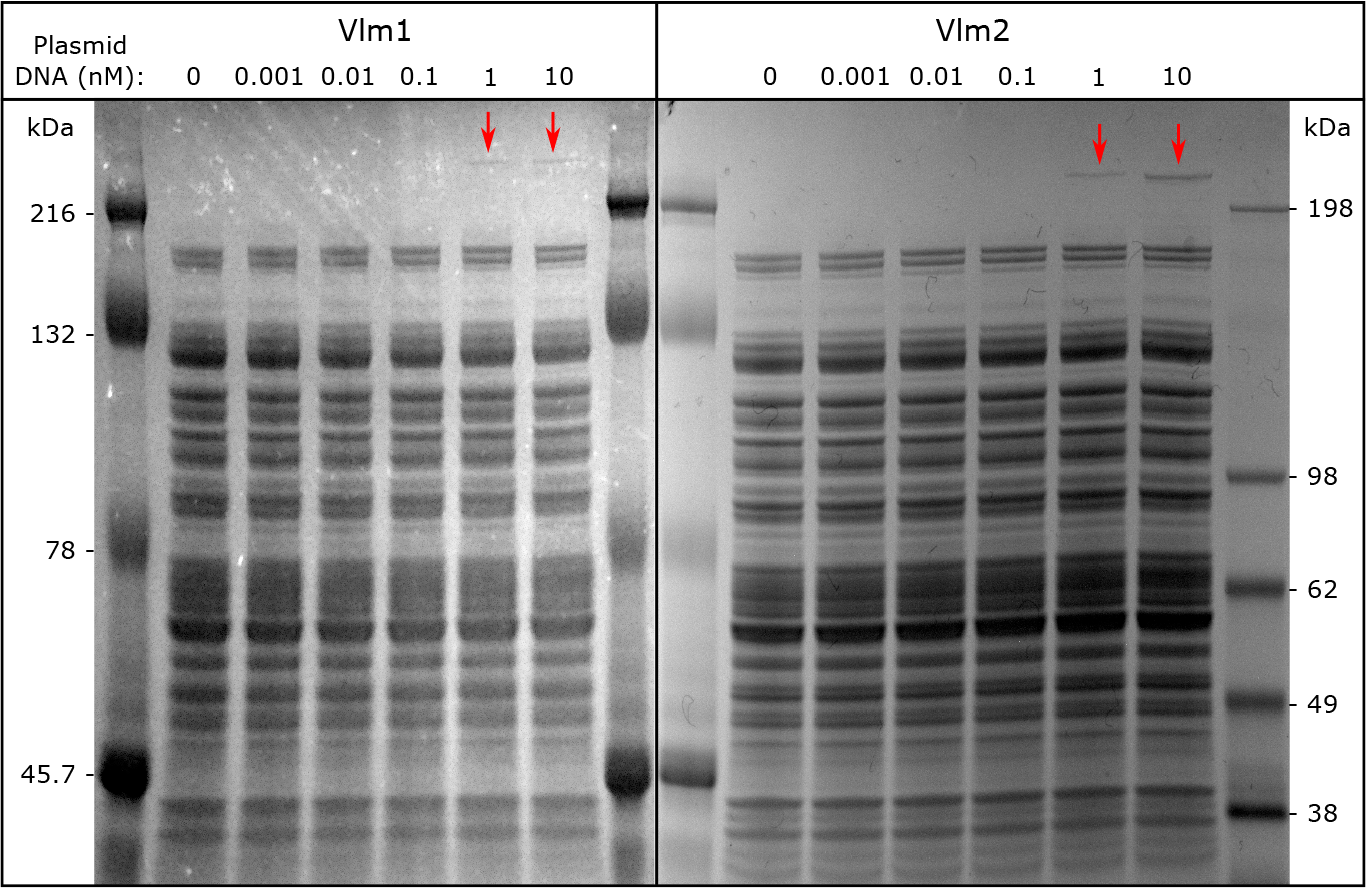
SDS-PAGE gels for Vlm1 (left) and Vlm2 (right) expression in TX-TL extract made from *E. coli* strain BJJ01. Different concentrations of pVlm1 and pVlm2 plasmid DNA were tested. Expected sizes are 370kDa for Vlm1 and 284kDa for Vlm2. Bands indicating successful expression of Vlm1 and Vlm2 appear at 1nM and 10nM concentrations (marked by arrows).

**Figure 5:**
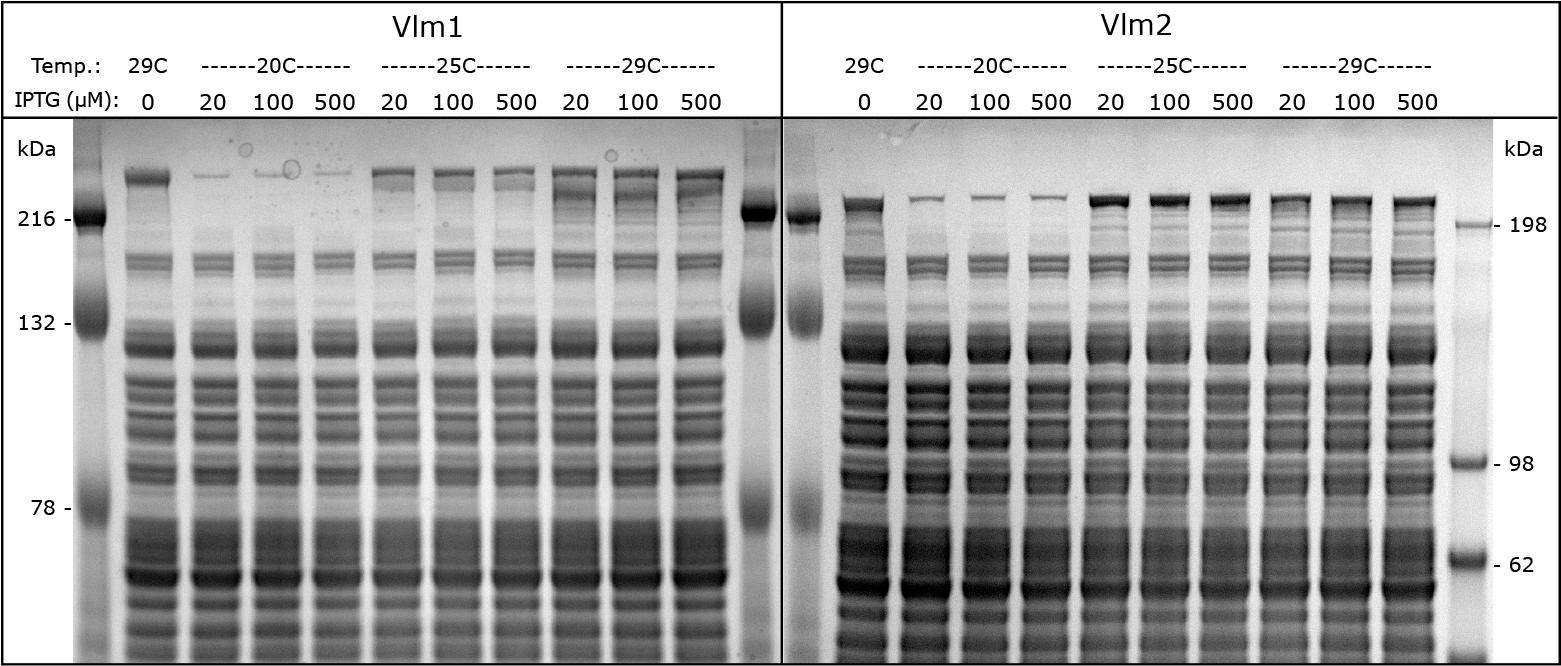
SDS-PAGE gels for Vlm1 (left) and Vlm2 (right) expression in TX-TL extract made from *E. coli* strain BJJ01. pVlm1 and pVlm2 plasmid DNA concentrations were kept constant at 1nM, while reaction temperature and IPTG concentration were varied. Expected sizes are 370kDa for Vlm1 and 284kDa for Vlm2.

### 3.4 Testing a fluorescence assay for valinomycin in TX-TL

Currently, the most reliable quantitative assay for detecting small molecules like valinomycin is LC-MS, but it requires expensive equipment and has only moderate throughput. Therefore, we investigated an alternative assay using the fluorescent probe 1-anilino-8-naphthalene sulfonate (ANS). Valinomycin is an ionophorous antibiotic, meaning that it forms positively charged complexes with alkali-cations, such as K+, Na+, and Rb+, to affect ion transport in bacteria [15]. When a cation binds to valinomycin, a conformational change occurs, creating a hydrophobic cavity which ANS can detect.

Feinstein & Felsenfeld (1971) were able to determine fluorescence emission as a function of valinomycin, cation, and ANS concentration in water [15]. We performed a similar experiment by combining different concentrations of pure, analytical grade valinomycin, KCl, and ANS both in water and in TX-TL, expecting fluorescence to increase with valinomycin. However, we found only slight increases in fluorescence intensity when in water, and no increases when in TX-TL (Figure 6). Further tests are needed, as we may not have used the optimal excitation/emission wavelengths in these measurements. If successful, this assay will provide a faster, easier way to quantify valinomycin production.

**Figure 6:**
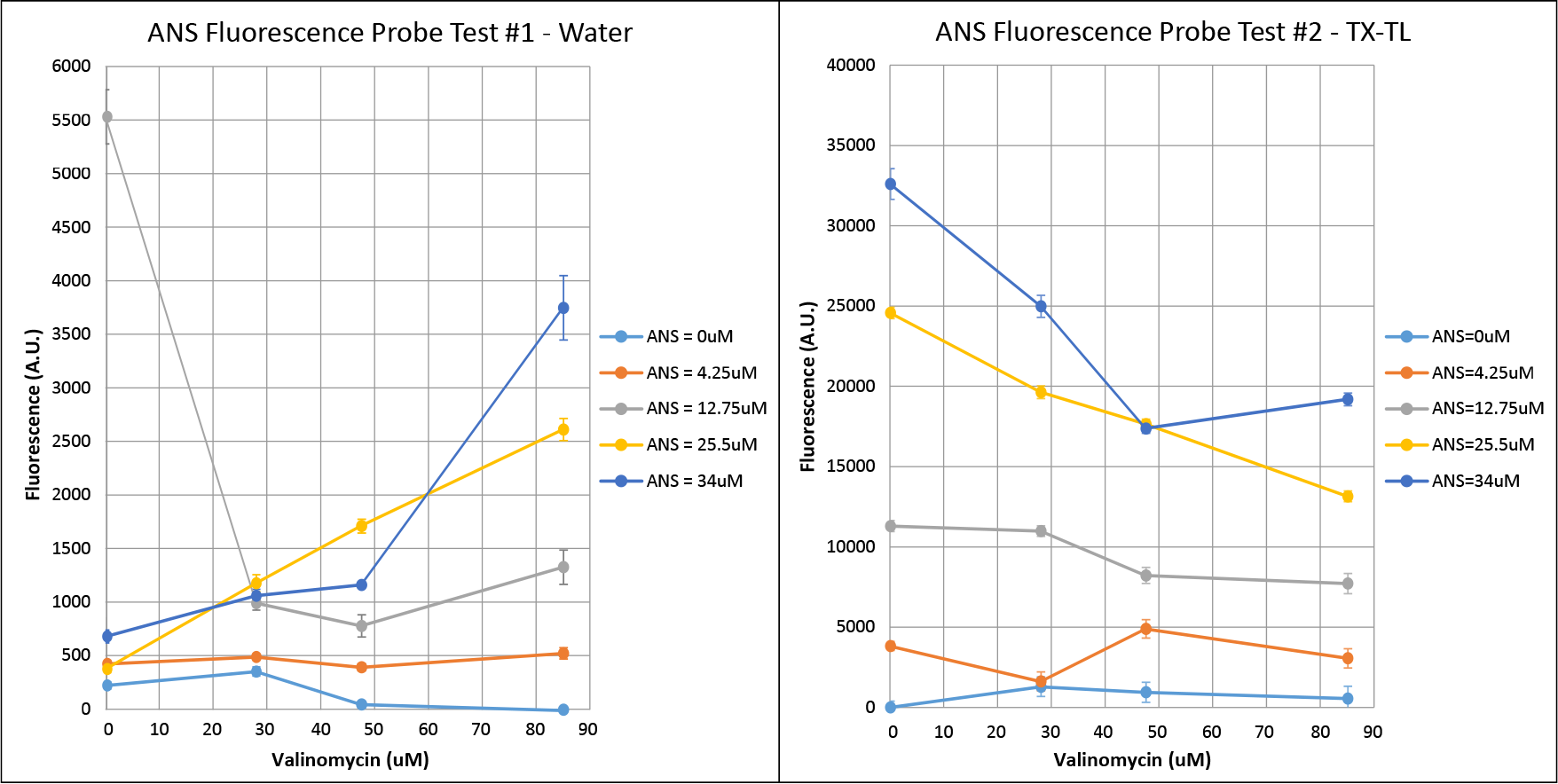
Results of ANS fluorescence probe test in water (top) and in TX-TL extract made from *E. coli* strain BJJ01 (bottom). Different concentrations of ANS and pure valinomycin were combined, along with KCl at a constant concentration of 225mM in both tests. Fluorescence was measured using 380/500nm excitation/emission wavelengths at gain 100.

### 3.5 Using LC-MS to detect valinomycin in TX-TL

Since the ANS fluorescence assay was unsuccessful, we moved on with LC-MS analysis. We added pVlm1 and pVlm2 together into a TX-TL reaction, along with the precursor L-Val in purified form. We also added purified D-Val because, at the time, we mistakenly thought it was a required precursor. We did not feed in *α*-ketoisovalerate (*α*-Kiv) and pyruvate (Pyr) because we were unable to obtain purified solutions for them at the time of the experiment. After running the TX-TL reactions at 25°C for 12 hours, they were analyzed using LC-MS, with a purified 0.5mg/L valinomycin standard as a positive control. The ion chromatogram (Figure 7) shows that the case of 1nM pVlm1 and 0.5nM pVlm2 results in the best production, about 0.6mg/L (Table S2, Supplementary Material). Since the TX-TL samples were diluted 20x prior to LC-MS analysis, the actual amount of valinomycin produced in this case is about 12mg/L. Adding less than 1nM of pVlm1 leads to decreased valinomycin production, even when pVlm2 concentrations are higher, suggesting that Vlm1 expression needs to be higher than Vlm2 expression. Adding 1nM of each plasmid decreases valinomycin production as well, possibly due to resource limitations preventing optimal Vlm1 and Vlm2 expression. In the MS spectrum (Figure 8), we see mostly the ammoniated adduct of valinomycin, analogous to the results in the Jaitzig et al. paper [9], confirming that we are seeing the correct product.

**Figure 7:**
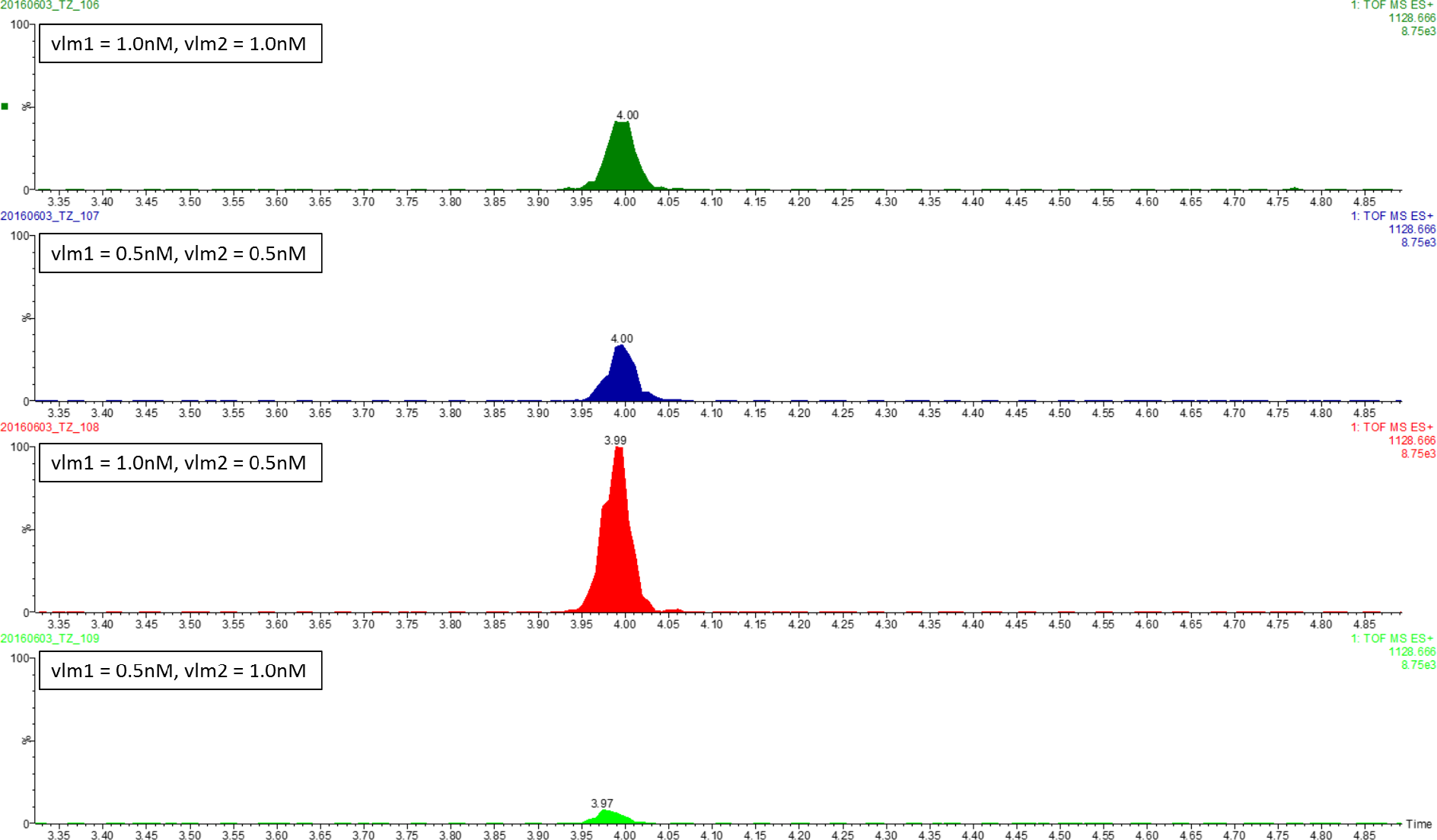
Ion chromatograms from LC-MS analysis of valinomycin production in TX-TL extract made from *E. coli* strain BJJ01. Not shown: Positive control has a peak similar to the “vlm1 = 0.5nM, vlm2 = 0.5nM” case, while the negative control has no visible peak.

**Figure 8:**
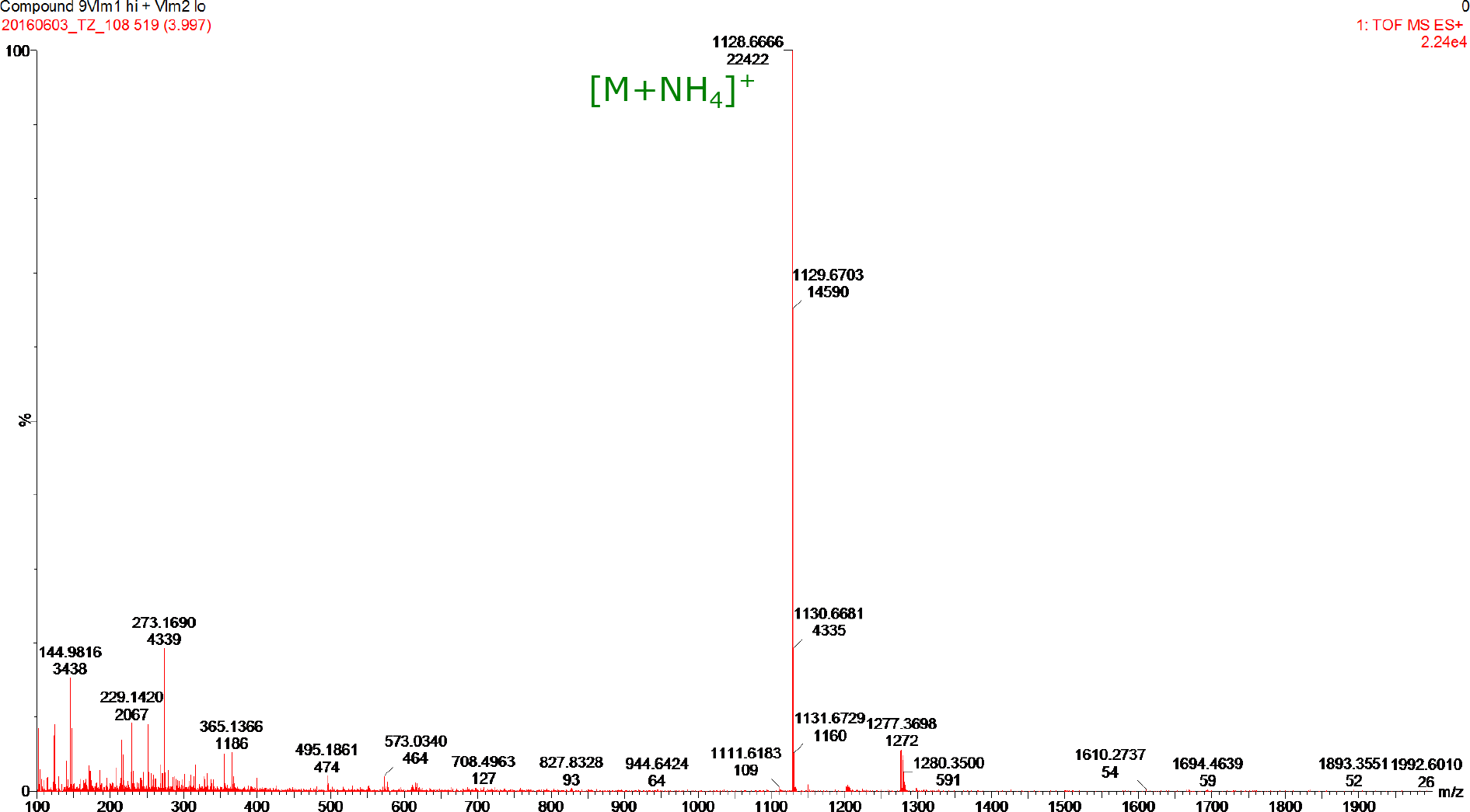
MS spectrum of valinomycin peak for the best-producing “vlm1 = 1nM, vlm2 = 0.5nM” test case. We mostly see the ammoniated adduct of valinomycin ([M+NH4]+ *m/z* 1128.6), similar to what was seen in the Jaitzig et al. paper [9].

## 4 Conclusion

Our results reveal that the pathway for valinomycin biosynthesis (originally from *Streptomyces tsusimaensis*) is able to function within a heterologous *E. coli*-based cell-free TX-TL system. Using extract created from *E. coli* BJJ01, which constitutively produces Sfp (needed for post-translational modifications to the VlmSyn proteins), we were able to detect valinomycin production of up to 12mg/L after the addition of plasmids containing the *vlm1* and *vlm2* genes. Only one out of the three precursors for valinomycin biosynthesis, L-Valine, was fed into the system, suggesting that the other two basic precursors, *α*-ketoisovalerate and pyruvate, must already be present in the extract. In the future, more concentration combinations of the plasmids will be tested, along with linear DNA versions of the genes, which may reduce resource loading and increase yield. Additional tests should be run to determine optimal precursor feeding concentrations, or whether precursor feeding is even required. Finally, a faster, easier assay for valinomycin, such as the ANS fluorescent probe, should be further developed in order to speed up testing of the pathway.

## 5 Materials/Methods

### 5.1 Obtaining *vlm1* and *vlm2* genes

The plasmids used in this study are listed in Table S1 (Supplementary Material). PCR amplification of *vlm1* and *vlm2* was carried out using Phusion Hot Start Flex 2X MasterMix (New England Biolabs) according to the manufacturer’s protocol with the following modifications: number of cycles increased to 40, extension time increased to 1 min/kb, annealing time increased to 1 min, initial denaturation time lengthened to 2 min, extension temperature increased to 75°C, and annealing temperatures varied between 68°C and 75°C on a gradient. Primers for PCR are listed in Table S3 (Supplementary Material) and were ordered from Integrated DNA Technologies. PCR products were analyzed on a 1% agarose gel.

### 5.2 Creating TX-TL extract from *E. coli* BJJ01

The strain *E. coli* BJJ01 was obtained from Jaitzig, et al. [9] as a glycerol stock. A small sample was streaked onto LB agar (no antibiotic) and incubated overnight at 37°C prior to extract preparation. The extract was made following the methods described by Sun, et al. [14], with the exception that we used homogenization instead of bead-beating during the cell lysis step. Buffer for this extract required 4mM of Mg-glutamate and 280mM of K-glutamate.

### 5.3 Expressing Vlm1 and Vlm2 in TX-TL

TX-TL reactions were conducted following the protocols described in [14]. When possible, DNA and inducers (such as IPTG) that were to remain at constant concentrations were added to a mastermix of extract and buffer to ensure uniform distribution. TX-TL reactions were conducted in PCR tubes at the standard volume of 10uL per reaction and incubated for 12 hours in a PCR machine. To prepare samples for SDS-PAGE, we mixed 1uL of the TX-TL reaction sample with 10uL of 1X LDS buffer and then heated the mixture at 70°C for 10 min. Samples were loaded into pre-made Bolt 4-12% Bis-Tris Plus Gels (Thermo Fisher Scientific Inc.) and run at 200V for 30-40 min. Gels were stained with SimplyBlue SafeStain (Thermo Fisher Scientific Inc.) following the manufacturer’s protocol.

### 5.4 Testing a fluorescence assay for valinomycin in TX-TL

The fluorescent probe 1-anilino-8-naphthalene sulfonate (ANS) was ordered from Sigma-Aldrich (Product Number A1028), along with ready-made valinomycin standard (~1 mg/mL in DMSO, 0.2 um filtered, Product Number V3639). Initial stock solutions of ANS and valinomycin were obtained by diluting with water; different concentrations were then obtained from these stocks by further diluting with either water or TX-TL depending on the experiment. Total volume per reaction was 10uL. Reactions were conducted in a 384-well plate, and fluorescence was measured at 29°C using a plate reader with 380/500nm excitation/emission wavelengths at gain 100. Three consecutive measurements were taken for each reaction.

### 5.5 Using LC-MS to detect valinomycin in TX-TL

We obtained L-Valine (reagent grade, ≥98%, Product Number V0500) and D-Valine (≥98%, Product Number 855987) from Sigma-Aldrich. TX-TL reactions were conducted following the protocols described in [14], in PCR tubes at the standard volume of 10uL per reaction, and incubated at 25°C for 12 hours in a PCR machine. Ten repeats were made for each reaction in order to have enough volume for LC-MS analysis. After completion of the TX-TL run, the repeats were combined and diluted 1:20 with water. LC-MS analysis of extracts was performed with the Waters Acquity UPLC-TOF MS system. For UPLC separation, we used a 2.1 *×* 50 mm, 1.8 um column with water +0.1% HCOOH (v/v) as eluent A and acetonitrile +0.1% HCOOH (v/v) as eluent B. Samples were separated at a flow rate of 0.3 mL/min with a linear gradient from 5 to 100% B over 2.5 min, an isocratic elution at 100% B for 5.5 min and a linear gradient from 100 to 5% B over 4 min. In order to quantify valinomycin produced in the TX-TL reactions, a purified 0.5mg/L valinomycin standard was used as a basis for comparison.

## 6 Acknowledgments

We would like to thank Prof. Dr. Peter Neubauer (TU Berlin) and Dr. Jian Li (Northwestern University) for providing us with their pVlm1, pVlm2, and sfp plasmids, their *E. coli* BJJ01 strain, and their helpful comments regarding the valinomycin biosynthesis pathway. We are also thankful for Dr. Nathan Dalleska (Caltech) for facilitating the LC-MS analysis, Yong Wu (Caltech) and Clare Hayes (Caltech) for providing input on techniques used in lab, and Dr. Richard Murray (Caltech) for providing valuable advice throughout the course of the project and the permission to conduct research in his laboratory. Funding for this project was provided by the Caltech Grubstake program.

## Supplementary Material

**Table S1:** Plasmids used in this study.

| Plasmid | Description | Source |
| --- | --- | --- |
| pVlm1 | *plac_CTU* promoter, SD_T7 RBS, N-terminal His-tag, attB sites, *lacI*, pMB1 ori, Cm<sup>R</sup>, *vlm1* | Jaitzig et al.[9] |
| pVlm2 | *plac_CTU* promoter, SD_T7 RBS, N-terminal His-tag, attB sites, *lacI*, parB locus, pMB1 ori, Amp<sup>R</sup>, *vlm2* | Jaitzig et al.[9] |
| pET15b-sfp | *pT7* promoter, *His-sfp*, *lacI*, *rop*, T7 terminator, pMB1 ori | Jaitzig et al.[9] |

**Table S2:**
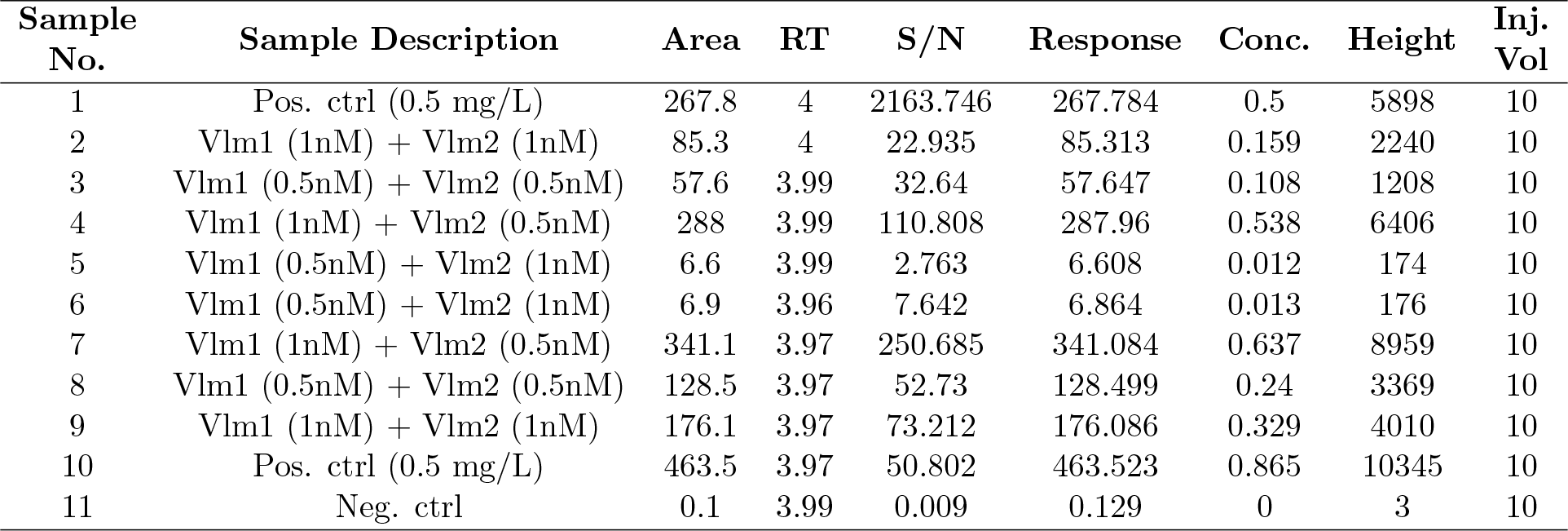
LC-MS results.

**Table S3:**
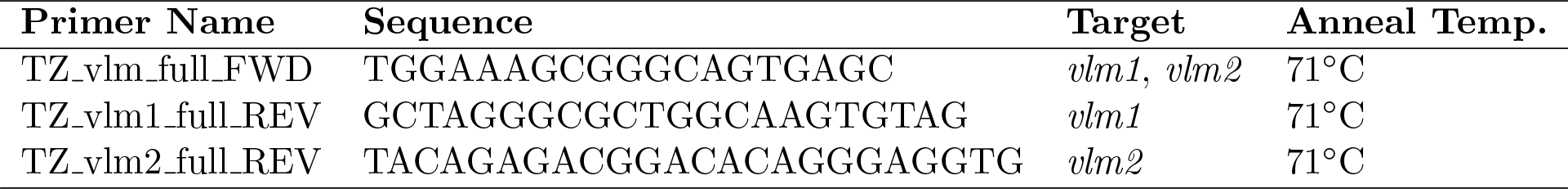
Primers used in this study.

